# Non-equilibrium vibrational dynamics govern ultrafast electron transfer in avian cryptochrome

**DOI:** 10.64898/2026.05.25.727693

**Authors:** Jonathan Hungerland, Daniel Timmer, Anders Frederiksen, Daniel C. Lünemann, Dustin Thöle, Ghazaleh Saberamoli, Jessica Schmidt, Krishan Kumar, Rabea Bartölke, Antonietta De Sio, Henrik Mouritsen, Christoph Lienau, Ilia A. Solov’yov

**Affiliations:** Institute of Physics, Carl von Ossietzky Universität Oldenburg, Carl-von-Ossietzky-Straße 9-11, 26129 Oldenburg, Germany; Department of Chemistry, School of Engineering Sciences in Chemistry, Biotechnology and Health (CBH), KTH Royal Institute of Technology, Stockholm, Sweden; Institute of Biology and Environmental Sciences, Carl von Ossietzky Universität Oldenburg, Carl-von-Ossietzky-Straße 9-11, 26129 Oldenburg, Germany; Research Centre for Neurosensory Science, Carl von Ossietzky Universität Oldenburg, Carl-von-Ossietzky-Straße 9-11, 26129 Oldenburg, Germany; Center for Nanoscale Dynamics (CENAD), Carl von Ossietzky Universität Oldenburg, Carl-von-Ossietzky-Straße 9-11, 26129 Oldenburg, Germany

## Abstract

Photoactivated intermolecular electron transfer (ET) in cryptochromes proceeds along chains of aromatic residues and creates a spatially separated pair of radical electrons. Ultrafast time-dependent spectroscopy can provide experimental insight into this process and theoretical estimates of charge transfer rates are commonly obtained via Marcus theory. Here, we present a new perspective on the ET in *European robin* cryptochrome 4a (*Er*Cry4a) that synthesizes insights from real-time ET calculations, ultrafast spectroscopic measurements and analytical derivations. The simulations exemplify that molecular vibrations play an essential role in enabling the ET dynamics, which was further rationalized through analytical derivations. Ultrafast pump–probe spectroscopy provided experimental access to the first 1.5 ns of the ET cascade, where multiple radical pair recombination rates arise due to the dynamic equilibrium along the ET chain. We show that the motions of the protein environment and the ET dynamics are inseparably coupled, violating the timescale separation required for Marcus theory. The presented results highlight that non-equilibrium coupling between electronic and nuclear motion dominates ET kinetics in *Er*Cry4a during the first nanosecond after photo-excitation. The findings exemplify the limits of Marcus theory and refine the interpretation of ultrafast spectroscopic signatures in cryptochromes.

## Introduction

Intermolecular electron transfer (ET) is of central mechanistic relevance for many biological systems. Within the respiratory chain [1] and photosynthetic complexes [2], ET chains within the proteins enable the conversion of chemical energy into an electrostatic gradient across a membrane and in photolyase proteins, DNA repair relies on ETs [3–5]. In the present manuscript, the protein cryptochrome 4a from *European robin* (*Er*Cry4a) is investigated, which is thought to be involved in magnetoreception [6–13], due to a spin-correlated pair of radical electrons that can be formed within the protein [10]. The radical pair is created by a chain of ETs within *Er*Cry4a, that occur after the non-covalently bound chromophore flavin adenine dinucleotide (FAD) has been excited by blue light (see Fig. 1a) [6, 14]. Accumulated evidence from bird genomes [15], gene expression patterns [16, 17], localization of protein expression [8, 18] as well as behavioral experiments involving light-dependence [19] and radio frequency noise [20–25] support the rationale that *Er*Cry4a is the sensory protein for the radical pair based magnetoreception process in birds. The potential signaling mechanism relies on the influence of the geomagnetic field on the spin dynamics of the radical pair on a timescale of 700 ns, which results in a shift of the singlet versus triplet radical pair populations [10, 21]. Simultaneously, relaxation and decoherence effects in the nanosecond regime disrupt the information content of the initially correlated radical pair [26, 27]. The radical pair created by the ET is the central sensory system as the spin-dynamics of the radical pair are modulated by the magnetic field [10, 21]. This modulation allows the weak magnetic field to shift the balance between a yet unknown spin-insensitive downstream reaction (but see [28–30]) and the singlet state specific recombination. An understanding of the ET process within *Er*Cry4a is therefore crucial to decipher or dispute the radical pair mechanism for avian magnetoreception.

**Figure 1:**
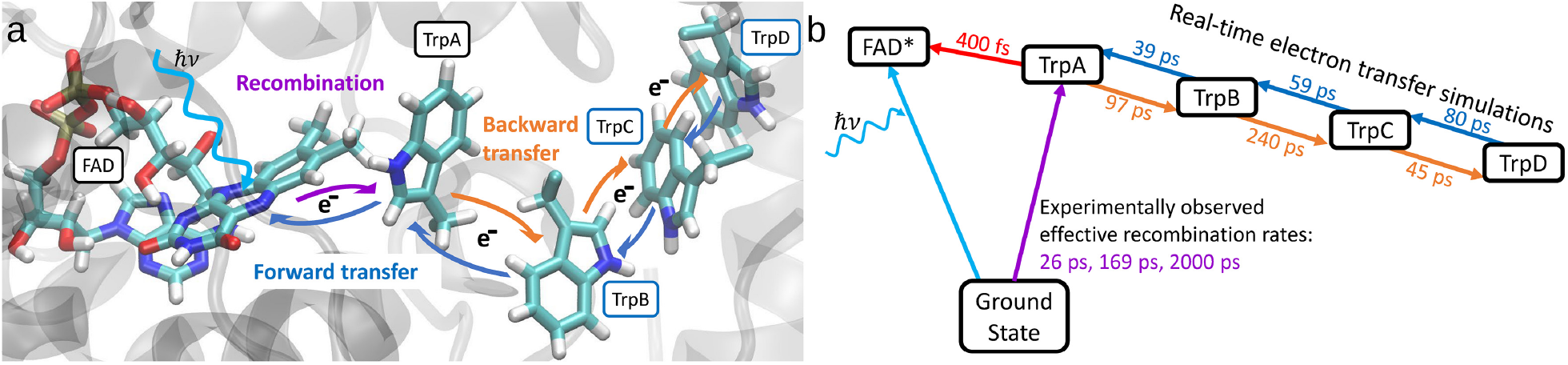
ET process in *Er*Cry4a. Dark-blue and orange arrows indicate the flow direction of electrons. **a:** Rendering of the non-covalently bound flavin adenine dinucleotide (FAD) and a chain of the four proximal tryptophan residues (TrpA-D) which constitute the ET transport chain. **b:** Overview of expected charge transfer transitions in *Er*Cry4a with rate constants taken from experiments (purple and red numbers) and simulations (blue and orange numbers). TrpX (X=A, B, C, D) abbreviates the radical pair state [FAD^•−^, TrpX^•+^] (see text). Light-blue, red and purple arrows indicate transitions between spectroscopically distinguishable states. The transfer times in b is a summary of the results of the present paper.

Figure 1 illustrates the dominant ET chain in *Er*Cry4a. The ET chain is initiated by the excitation of flavin which creates an electron hole in the lowest unoccupied molecular orbital that is filled by an outer shell electron of the proximal TrpA residue via the first *forward* ET. The ETs in the forward direction follow the expected free energy gradient. Polar and charged moieties of the protein like TrpH^•+^ interact favorably with the hydrophilic protein exterior, such that the ETs are expected to relocate the radical electron hole to the periphery of the protein. During a *forward* ET, for example 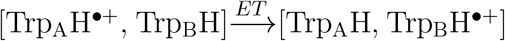 (denoted as *A*→*B*), the electron transfer to the protein interior brings the charged TrpH^•+^ closer to the protein exterior. Conversely, the ET process 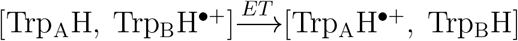 (denoted as *B*→*A*) is defined as a *backward* process. The final backward process 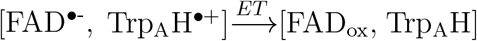 (denoted as *A*→*G*) is an irreversible charge *recombination* as it returns the protein into a closed-shell ground state configuration by recombining two radical electrons.

Here, we provide multiple perspectives to elucidate the ET events in *Er*Cry4a within the first 1 ns after photo-excitation. We suggest a model of vibrationally assisted electron transfer that consistently synergises computational and experimental data. The model agrees well with experimental observations on similar flavoproteins [31–34] and with theoretical expectations on the coupling between ET and molecular motion [35–38]. On the example of photo-activated *Er*Cry4a, we dissect how vibrations, ET and molecular motion are coupled, why these couplings lead to a break-down of typically employed methodologies and where the ET-vibrational couplings originate. Additionally, we show how such a truly non-equilibrium ET process can be studied *in silico* and how this leads to a much better agreement between computational and experimental data and thereby a much improved understanding of the fundamental cryptochrome activation mechanism.

## Results

### Non-equilibrium breakdown of Marcus theory for ET rate estimation

Theoretical estimates of the ET rate constants in biological systems often rely on Marcus theory [39]. In a semi-classical formulation, the resulting charge transfer rate between two molecular sites reflects quantum mechanical wavefunction overlap as well as the donor–acceptor energy spectrum, while environmental interactions are treated classically. Such an approach allows to establish the charge transfer rate constant as [5, 39, 40]:

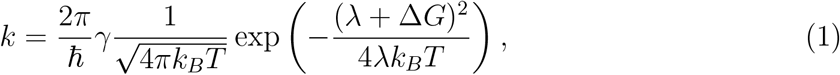

where *ħ* is the reduced Planck’s constant, *k*_*B*_ is the Boltzmann constant and *T* is the temperature of the system. The reorganization energy *λ* measures the energy needed to instantaneously change the electronic configuration of the protein into the post-transfer configuration, while the reaction free energy difference Δ*G* measures the equilibrium thermodynamic energy difference between the electronic states before and after the ET. The electronic coupling *γ* is a geometric quantity. It measures the overlap between the electronic wavefunction before and after the ET. For intermolecular ET, *γ* decreases approximately exponentially with increasing edge-to-edge distance between the molecular sites involved [41].

**Table 1:** Overview of computationally derived time constants for ET processes within *Er*Cry4a. For the *Full vibrations* column, real-time ET in a molecular tight-binding/AMBER force field embedding was conducted with free bond vibrations. For comparison, data from a previous study by Timmer *et. al*. [14], which constrained molecular vibrations in an otherwise identical setup, is reanalyzed. All other methods used molecular dynamics to sample various states of the ET chain and computed energies and electronic overlaps using embedded density functional theory (ΔSCF *ω*B97X-D/6-31G(d)/AMOEBA by Cascone *et al*. [41], TD-DFT *ω*B97X-D/6-31+G^**^/CHARMM by Liu *et al*. [6], CAM-B3LYP/6-31G/CHARMM by Xu *et. al* [11]), which were then interpreted via Marcus theory. For the ground state recombination process calculations, independent relative errors of *±* 5% are considered for every rate if no explicit error estimate is given in this table. Note, that *G* → *A* only occurs after photoexcitation of FAD.

| Step | Full vibrations | Timmer *et al.*<br>constrained vibrations | Cascone *et al.* | Luo *et al.* | Xu *et al.* |
| --- | --- | --- | --- | --- | --- |
| $G \rightarrow A$ | - | - | 830 fs | - | - |
| $A \rightarrow G$ | - | - | - | - | - |
| $A \rightarrow B$ | 39 ps | 20 ps | 15 ps | - | 1.1-1.8 ps |
| $B \rightarrow A$ | 97 ps | 82 ps | 32 ns | - | 56-125 ns |
| $B \rightarrow C$ | 59 ps | 6.4 ps | 77 ps | - | 15-31 ps |
| $C \rightarrow B$ | 240 ps | 0.35 ps | 2700 μs | - | 0.76-2.0 μs |
| $C \rightarrow D$ | 80 ps | 4.4 ps | 137 ps | 33 ps | 59-110 ps |
| $D \rightarrow C$ | 45 ps | 39 ps | 26 ns | 33 ps | 53-91 ps |
| $C \rightarrow G$ | - | - | - | 830 ns | 59-140 ns |
| $D \rightarrow G$ | - | - | - | 3.3 μs-33 ms | 2.0-5.3 μs |

Estimates of the ET in *Er*Cry4a based on Marcus theory have been previously carried out [6, 11, 41], their results are compiled in Tab. 1. The Marcus theory estimates differ primarily by the employed level of theory, the force-field used for the classical embedding model, sampling times and initial guesses of the protein models. Both sets of estimates predict forward electron transfer rates of 1-140 ps, while the back-transfer rates are orders of magnitude slower with typical time constants on scales from nanoseconds up to milliseconds. Following this order of magnitude difference in rates, the electron back-transfers towards the TrpB and TrpA sites are collectively identified as very unlikely [11, 41].

The Marcus theory approach typically involves classical, nanosecond-long molecular dynamics simulations of the protein, during which the structure is restrained to a particular radical pair state. The long dynamics allow the local [42] and global [43] protein structure to adapt to the particular location of the TrpH^•+^ species. The sampling procedure roots upon the ergodian sampling equilibrium assumption underlying Marcus theory, which considers the protein structure to be a sample from an equilibrium distribution of conformations. However, given the picosecond-scale ET rates, the protein structure relaxes too slowly and cannot be assumed to be in equilibrium during the ET process. The studied protein, therefore, is expected to demonstrate the non-Gaussian and non-ergodic nature of electron transfers in proteins [36–38].

### Beyond Marcus theory: real-time ET simulations

To account for the coupled dynamics of the electron hole and the protein environment, realtime quantum mechanical/molecular mechanical (QM/MM) calculations were performed. In this approach, an electrostatically embedded ET chain is treated quantum mechanically, while the rest of the system is described classically. In a previous study [14], an implementation that combines tight-binding density functional theory and all-atom molecular dynamics was utilized [14, 44]. This tight-binding description [44–48] allowed us to model the charge transfer across the tryptophans. Transfer events involving the flavin moiety were not considered and (de-)protonation events, bond-formations or bond-breaks were not modeled either. Additionally, the previous study [14] constrained the bond lengths between atoms in the protein and backward transfer processes were assumed to be negligible for the analysis of the rate constants. The present study refines the earlier QM/MM calculations, while permitting for the more realistic bond vibrations and the possibility for the electron back transfers to take place. The initial state of the simulated system is fixed such that the radical electron is located at the Trp_A_H^•+^ residue, mathematically represented by the state vector 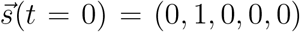. The time evolution of the average occupation of the state vector is then numerically modeled as

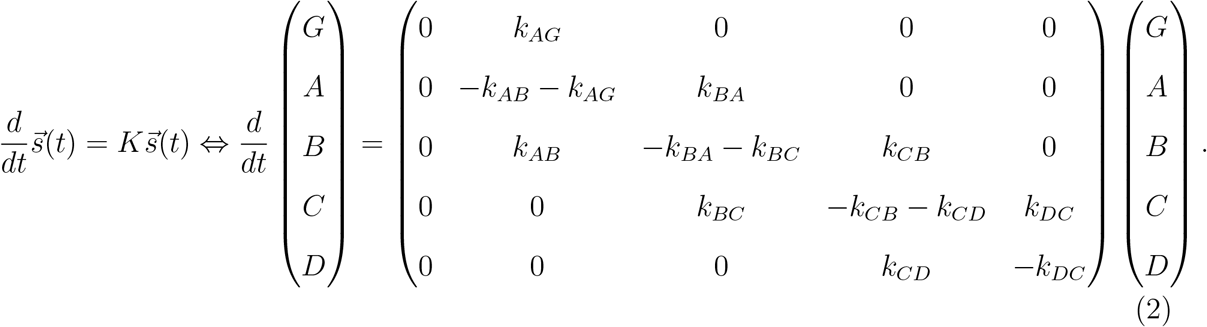

The coupling matrix *K* is fully defined by its strictly positive off-diagonal elements. The considered states are the ground state [FAD_ox_, TrpH] (*G*) and the radical pair states [FAD^•-^, Trp_X_H^•+^] (*X* = *A, B, C, D*). A transition from state *X* to state *Y* occurs with the rate *k*_*XY*_. The state vector represents a probability density, so ∑_*i*_ *s*_*i*_ = 1 always holds and the diagonal elements of *K* are thus directly expressed via the off-diagonal elements. Because recombination into the ground state *G* is considered irreversible, there is no rate *k*_*GA*_ in the coupling matrix *K*. In the ET simulations, the flavin moiety is not part of the QM region and the system starts in the *A* state. Thus, the final backward transfer *A*→ *G* can never be observed during simulations and fits to simulation data always yield *k*_*AG*_ = 0. Instead, *k*_*AG*_ will be inferred from experimental data [14].

Figure 2a illustrates the dynamics of the electron hole localization in the four Trp residues of *Er*Cry4a. Constrained vibrations cause the ET process to largely terminate at the TrpB residue within the studied simulation time, which is reflected in the rate constants shown in Tab. 1 through the fast backward-transfer *C*→*B*. In contrast, when realistic molecular vibrations are considered, the ETs occur generally faster and a dynamic equilibrium dominated by the *C* (0.54) and *D* (0.30) species is established within 300-500 ps after initial charge separation via *G*→*A*.

**Figure 2:**
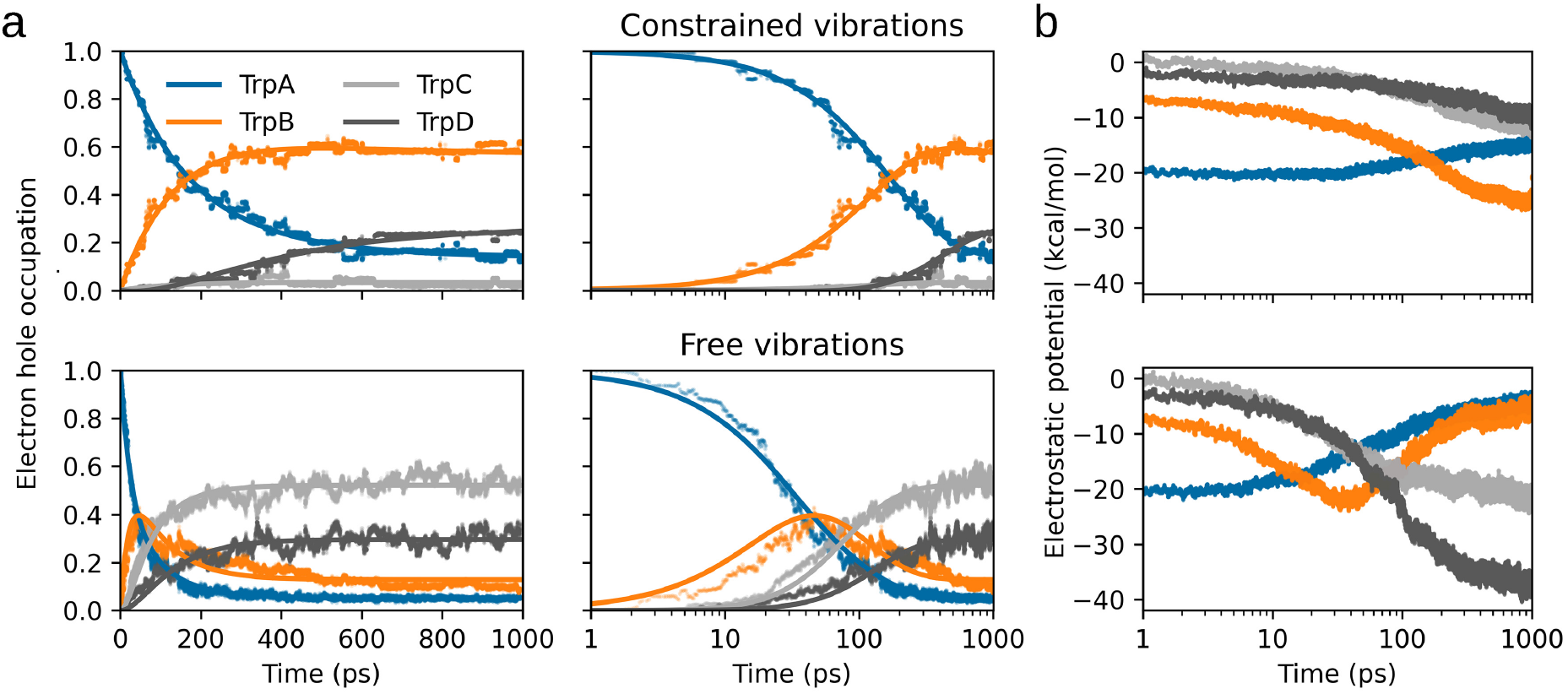
Real-time ET results with vibrational constraints (top row; data reanalyzed from [14]) and with free molecular vibrations (bottom row). **a:** Transfer of the radical pair measured as the occupation of an electron hole over time. An average over an ensemble of QM/MM trajectories is shown. The result of numerical fits including forward- and backward-transfer according to Eq. (2) are plotted as lines. **b:** Electrostatic potential energy of TrpA-D time-averaged over an ensemble of QM/MM trajectories.

Figure 2b shows the time evolution of the electrostatic potential of the possible radical pair states within *Er*Cry4a. A gradual change of the lowest-energy state from *A* to *B* within 10-30 ps is an effect of protein structural relaxation. This equilibration process establishes an electrostatic gradient that favors the forward charge transfer direction. Due to the structural rearrangements, the *C* state is energetically most favorable after 1 ns of simulation, even though the electrostatic potential appears 20 kcal/mol less favourable compared to the *A* state at t=0 ns. At every time instance, the electrostatic potential of the tryptophan residues is a good predictor of the currently most occupied state. The faster response of the protein environment to the moving charge in the case of realistic molecular vibrations allow the protein to adapt an equilibrium configuration more rapidly. Even though the electron transfer rates that follow from the kinetic model fitting, as listed in Tab. 1, are faster once molecular vibrations are constrained, the vibrationally assisted protein relaxation leads to the equilibrium configurations of the *C* and *D* states on a shorter time scale and manifests clearly within a time interval of 1 ns. Note, that in the case of vibrationally assisted electron transfer, one observes similar population dynamics after 30 ps, as in the case of simulations with constraints after 1 ns. This finding suggests that the charge transfer events could also be completed in the constrained simulations, at significantly longer timescales.

### The relevance of molecular vibrations for ET simulations

The differences between the sets of real-time ET transfer simulations due to the bond constraints are rationalized in Fig. 3a-b. In the constrained simulation, the bond-lengths are restrained to an average deviation of about 0.005 Å from their equilibrium value (see Fig. 3a). Such a restraint shifts the spectrum of bond vibrational frequencies by an order of magnitude and leads to a slight increase of angular vibrational frequencies (see Fig. 3b). Angular vibrations are those that involve at least two chemical bonds, which include, for example, bending, scissoring or twisting motions in the protein. Since the numerical restraints of bonded interactions allow only miniscule deviations in bond length, their effect is similar to increased force constants for the chemical bonds. These observations alone, however, are insufficient to predict the effect of vibrational constraints on the associated electron transfer rates.

**Figure 3:**
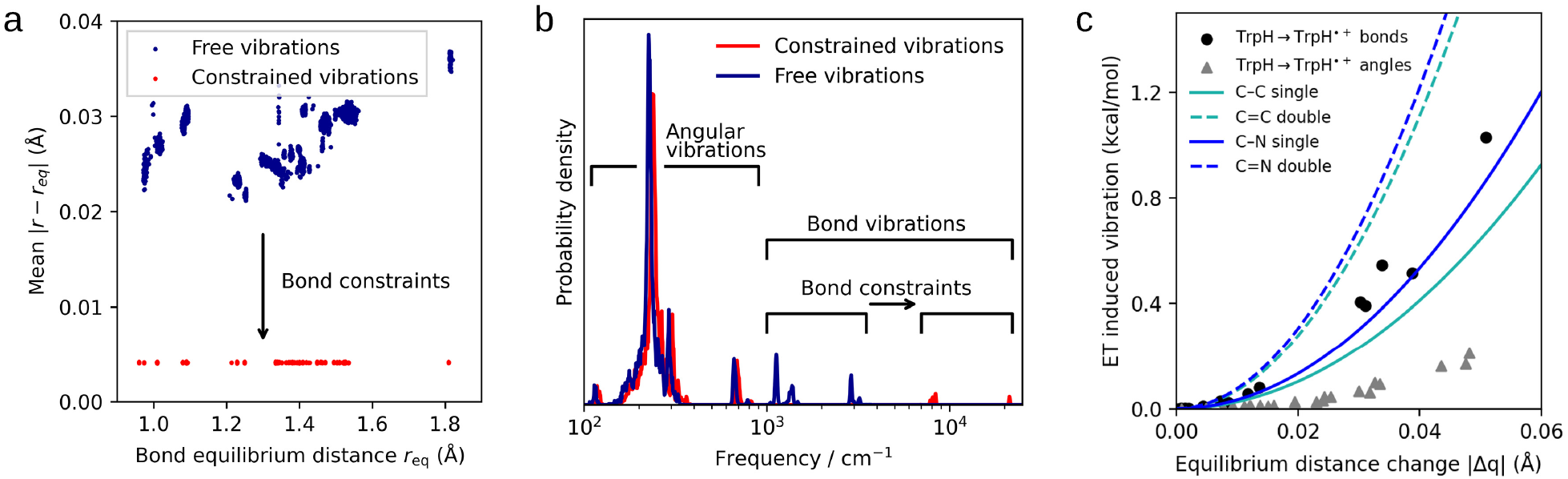
Effects of molecular bond constraints on vibrations and ETs. **a:** Average deviations from equilibrium bond lengths for the simulations with unconstrained bonds (blue) and constrained bonds (red) illustrate the procedure of numerical bond constraints. **b:** Effective vibrational frequency spectra highlighting the effective shift of bond force constants. **c:** Expected energy associated with vibrations and induced via ET for the TrpH → TrpH^•+^ process (symbols) compared with stretching modes of typical bond strengths (lines).

A complementing insight into the problem could be achieved through an extension of Marcus theory that accounts for the interplay between electron transfers and molecular vibrations [40, 49]. The extended theory accounts for the fact that upon ionization of a molecular moiety, e.g. when TrpH becomes TrpH^•+^ through ET, the energetic optimum of nuclear positions changes. For example, the optimal lengths of certain chemical bonds tend to shorten upon removal of an electron. Assuming that all molecular vibrations can be described via a collection of quantum harmonic oscillators and furthermore assuming that vibrational frequencies *ω* remain approximately identical before and after ET, one may arrive at an analytic expression of the vibrationally assisted ET rate constant. The full derivation of this expression will be discussed in detail in a more specialized journal. However, the relevant molecular Huang-Rhys factors are analysed here as these appear sufficient to explain the vibrational assist observed in the real-time QM/MM simulations. The molecular Huang-Rhys factor is a generalization based on the work of Solov’yov *et. al*. [49] that recovers the vibronic Huang-Rhys factor model [50] or the displaced harmonic oscillator model [51] in the limit of small coupling strengths (see supplementary results for further discussion).

Upon a charge transfer process among the tryptophan residues, equilibrium bond lengths within the residue changes by the distance *q*, which can also be expressed in terms of the dimensionless normal coordinate *u* defined as

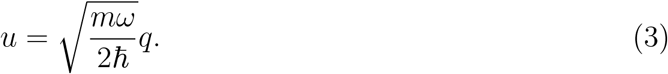

Here, *ω* is the frequency of the normal vibration and *m* is the effective mass of the involved atoms. Consider the mode to be initially in the state |*k*⟩, and after the ET, to populate state |*n*⟩. The change in an equilibrium distance within the tryptophan side chain can be expressed through the displacement operator 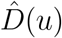, such that 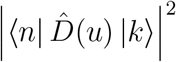, represents an overlap integral between the displaced and nondisplaced vibrational eigenfunctions. An analytical derivation [40] within the formalism of second quantization leads to

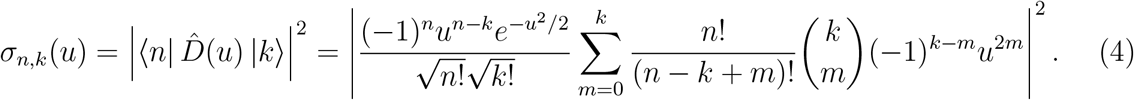

Here, the term *σ*_*n,k*_(*u*) is the molecular Huang-Rhys factor that defines the occupation of the vibrational state *n* by the tryptophan sidechain, given an initial vibrational state *k* and a shift of the equilibrium distance *u*. The displacement operator represents a repositioning of a wave function in real space, while the Huang-Rhys factor expresses how such a shift can be translated into a vibration. For example, consider a quantum harmonic oscillator in its vibrational ground state *k* = 0 that experiences the shift *u* = 1. Computing the Huang-Rhys factor according to Eq. (4) results in the following populations of the vibrational states after the shift for the states *n* = (0, 1, 2, 3, …):

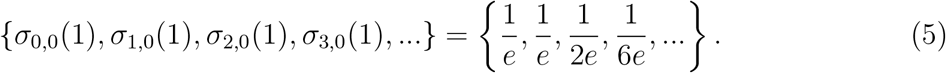

The estimate shows that after the shift, only *σ*_0,0_(1) = 1*/e* ≈ 37% of the wavefunction is still in a vibrational ground state, so the shift induced vibrations (*n >* 0 is occupied) for the previously non-vibrating (*k* = 0) system.

To further illustrate the meaning of the Huang-Rhys factor, a molecular system comprising the tryptophan sidechain is considered. Quantum chemical geometry optimizations and vibrational frequency calculations at the M06-2X/def2-TZVPD level of theory [52] were carried out using ORCA [53, 54] for the aromatic part of a tryptophan sidechain in its neutral form and positive radical form. The occupations *p*(*k*) of the vibrational modes *k* with frequencies *ω* are assumed to follow a Boltzmann distribution:

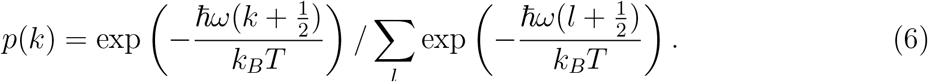

The expectation value of the vibrational energy after the ET could then be readily computed as

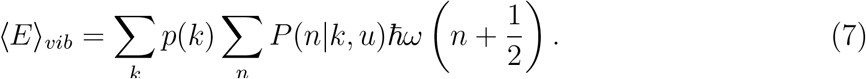

Here, the eigenvalue of the vibrational energy in mode *n* is *ħω*(*n* + 1*/*2). The term *P*(*n*|*k, u*) describes the probability to occupy a vibrational mode *n* given an initial vibrational mode *k* and an equilibrium distance shift *u*. This is exactly what the Huang-Rhys factor describes, i.e. *P*(*n*|*k, u*) = *σ*_*n,k*_(*u*).

The results from Eq. (7) for the tryptophan sidechain are shown in Fig. 3c. Absolute changes in bond lengths are given in Å, while the changes in angular internal coordinates are approximated as changes in distance, assuming that the bonds forming the angle remain of fixed length. Every point in Fig. 3c shows the contribution of one internal coordinate to the change in vibrational energy. The typical force constants for carbon-carbon bonds (1000 cm^-1^ single bond; 1650 cm^-1^ double bond) and carbon-nitrogen bonds (1100 cm^-1^ single bond; 1665 cm^-1^ double bond) were also used to illustrate the typical energy associated with stretching molecular bonds by a value of |Δ*q*|, see lines in Fig. 3c.

The estimates of the Huang-Rhys factor used to analyse the vibrational energies in Fig. 3c suggests that ETs induce molecular vibrations that are relevant on thermodynamic energy scales and effectively heat up the constituents of the ET chain (sum over bonded energies: 4.9 kcal/mol, sum over angular energies: 1.1 kcal/mol). During the ET process, electrostatic potential energy is converted into molecular vibrations. As long as the vibrational energy does not dissipate, the tryptophan moiety that hosts the radical electron is often found in an energetically degenerate state compared to any neighbouring tryptophans, which facilitates ET. The molecular vibrations, partially induced through ET, enable the transferring electron to hop more seamlessly in either direction of the transfer chain. In practical applications of the Marcus theory approximation, excited vibrational states are dissipated simply through the long timescales over which the structure was allowed to equilibrate. On the relaxation time-scales of global protein conformations (ns-*µ*s), Marcus theory estimates are expected to become appropriate [37].

### Implications for the interpretation of ultrafast spectroscopy data

If the general observation, that the sub-ns regime of ET processes is a dynamic, vibrationally excited non-equilibrium state, is true, then one must consider the implications for the interpretation of experimental data. Experimental access to ET processes has so far been gained via optical pump-probe spectroscopy utilizing narrowband, ultrashort pump pulses that are tuned into resonance with the sample absorption in combination with broadband probe pulses that cover all optical transitions of interest [14, 55]. A time-dependent differential transmission spectrum of the protein (see Fig. 4a), can be recorded systematically as a function of time relative to an excitation pulse at 450 nm, which corresponds to the peak of the flavin absorption band and, where flavin is pumped into an electronically excited state [56]. By comparing the excitation-induced transmission difference, a time-dependent differential transmission spectrum, ΔT/T, is obtained. ΔT/T quantifies the effect of the flavin’s excitation via the pump pulse on the recorded spectrum at different wavelengths over time. After extracting a selection of wavelengths (see Fig. 4c), one observes different time dependencies for a range of wavelengths, which suggests that a single Δ*T/T* spectrum that decays with some (multi-)exponential process is insufficient to explain the measured data. To account for this, global analysis [57] fits decay associated difference spectra (DADS) to the data as

**Figure 4:**
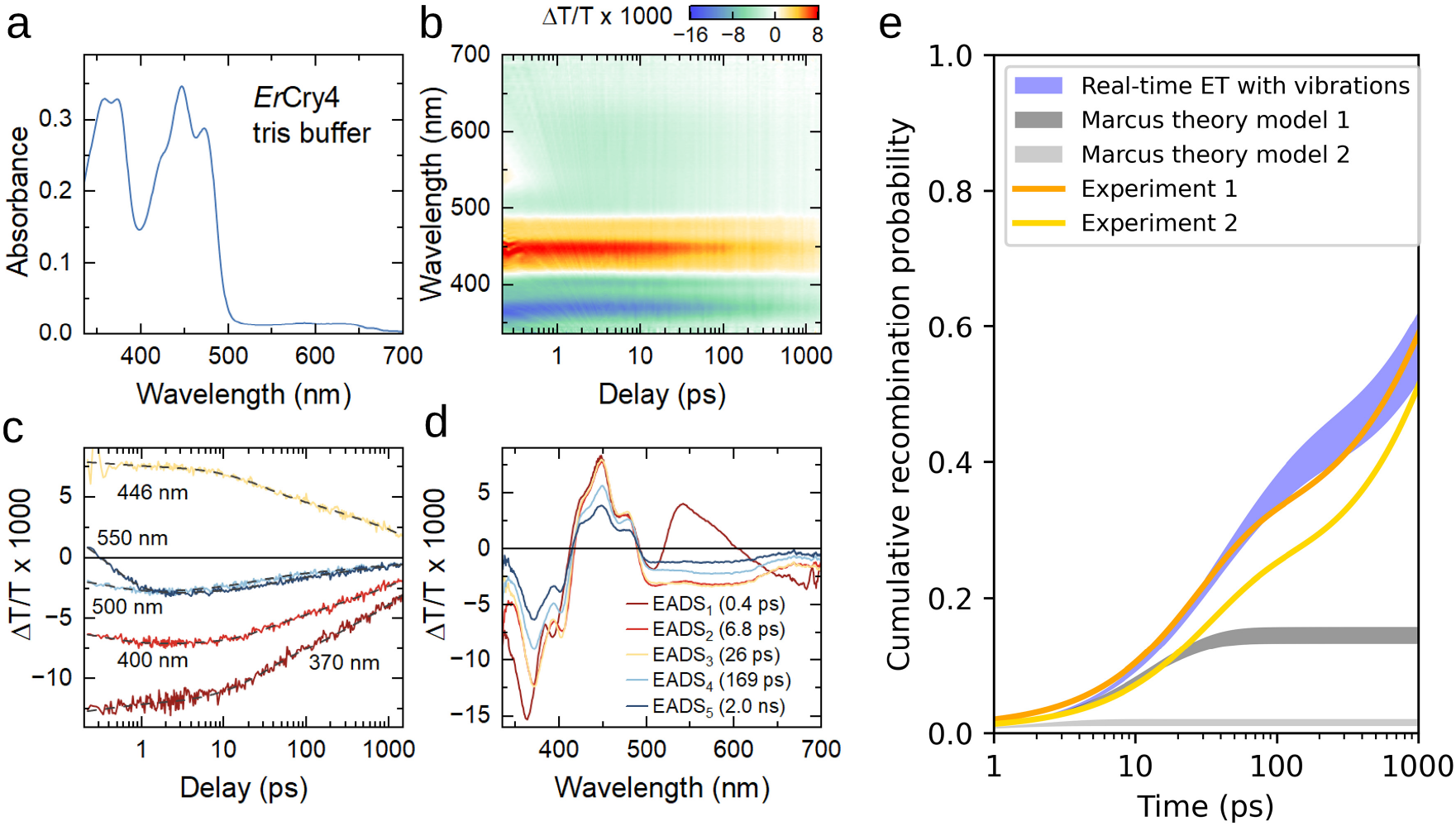
Experimental data for ET dynamics in wildtype *Er*Cry4a protein in tris buffer recorded at 1°C. **a:** Linear absorption spectrum of a 200 *µ*M sample of *Er*Cry4a. The main absorption band of FAD_ox_ is seen centered around 450 nm. Weaker absorption between 500-650 nm originates from small contribution of FADH present in the sample after photoreduction. For pump-probe, 2 mM potassium ferricyanide is added to reoxidize photoreduced flavins. **b:** Pump-probe map showing differential transmission spectra ΔT/T recorded up to a delay of 1.5 ns. A rapid decay of a positive stimulated emission signal close to 550 nm is the characteristic signature of the first ET from TrpA to FAD. **c:** Dynamics at selected wavelengths and fit resulting from the global analysis (dashed grey lines). **d:** Evolution-associated difference spectra (EADS) as obtained from global analysis of the data. **e:** Cumulative recombination probability over time as predicted via multi-state Markov models utilizing transfer rates from theoretical calculations compared with the experimentally obtained time-evolution.

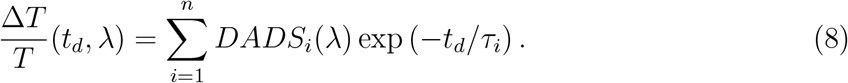

Each decay component is assigned its own characteristic absorption spectrum *DADS*_*i*_ and decay time *τ*_*i*_.The sums of all DADS spectra results in the first *so-called* evolution-associated difference spectrum (EADS)

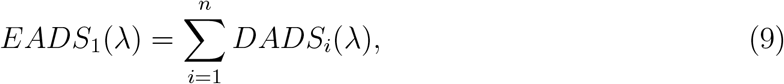

which contains all recorded spectral changes. Omitting some *DADS* components in the order of increasing *τ*, results in subsequent EADS spectra

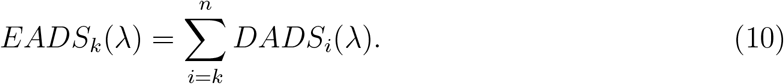

Thus, *EADS*_2_ contains all spectral decay that cannot be explained by the fastest decay component with decay time *τ*_1_. In other words *EADS*_2_ is the spectrum that evolves out of EADS_1_ over a characteristic time *τ*_1_. The EADS spectra measured for *Er*Cry4a are shown in Fig. 4d. Further in depth discussion of previous measurements and their spectroscopic assignment is given in an earlier study [14]. While those EADS components are rich in information, their interpretation must be done with care. *EADS*_1_ shows the spectral change associated with the formation of photoexcited FAD. *EADS*_1_ then decays with a time constant *τ*_1_=400 fs into *EADS*_2_. *EADS*_2−5_ feature the spectral shifts of the 362 nm and 391 nm bands caused by the change in redox state when forming the FAD^•−^ radical anion [14, 58]. Furthermore, the broad excited state absorption band replacing the stimulated emission signal in the spectral region between 500-650 nm is a characteristic signature for TrpH^•+^ formation [14, 33, 34]. Thus, *τ*_1_ can be assigned to the first ET that creates the radical pair state [FAD^•-^, Trp_A_H^•+^]. The evolution from *EADS*_2_ to *EADS*_3_ forms with *τ*_2_=6.8 ps and differs from *EADS*_2_ only by subtle changes in lineshape (Fig. 4d, red and yellow lines), such that it can be assigned to vibrational cooling dynamics [14, 33]. All following decays show comparable spectra associated with decays of [FAD^•-^, Trp_X_H^•+^] at varying time constants.

Experimentally, time constants of 29 ps, 169 ps and 2.0 ns were deduced for decay times *τ*_3−5_. The utilized optical markers for the radical pair formation, i.e. the absorption bands FAD^•-^ and TrpH^•+^ do not depend significantly on the location of the radical pair; thus, the states [FAD^•-^, Trp_X_H^•+^] (X=A,B,C,D) are spectroscopically indistinguishable. The decay of Δ*T/T* with time mainly reflects a decay of the total population of radical pairs. Multiexponential decays are observed, because with each forward ET recombination via *A*→*G* recombination becomes less likely, and the effective recombination rate slows down. The decay times can be a direct measurement of the forward ET rates but only if the backward rates are small compared to the forward transfer rates as previously assumed in Timmer *et. al*. [14]. Here, the forward and backward rates are similarly fast. Therefore, the decay times reflect the interplay between the recombination dynamics of the first radical pair via the *A*→*G* process and the dynamic equilibrium of forward and backward transfers along the ET chain.

The information inferred form the DADS analysis for the three slowest decay components can be combined into a multi-exponential decay model that quantifies the dynamics of the radical pair concentration. With *S*(*t*) being the population of radical pairs that survives after the first ET step and after vibrational cooling, the time evolution of *S*(*t*) can be expressed as

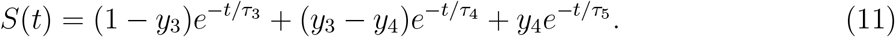

The above equation formalizes the analysis of the radical pair dynamics with the three rate constants *τ*_3−5_ = (26, 169, 2000) ps and four amplitudes *y*_2−5_ = (1.0, 0.71, 0.68, 0.0) deduced from the DADS analysis. The cumulative ground state recombination *G*(*t*) is computed as *G*(*t*) = 1 − *S*(*t*). Figure 4e shows *G*(*t*) for the experimentally obtained rate constants and yields.

The theoretical calculations deliver rate constants for the microscopic ETs along the tryptophan chain. In addition to those, the ground state recombination *A*→*G* needs to be included. Thus, the rate equation model in Eq. (2) now includes a non-zero rate *k*_*AG*_ for the recombination *A*→*G*, which was set to 89 ps following a result obtained for a mutated variant of *Er*Cry4a [14]. Only recombination from state A is included since recombinations from other states are expected to be orders of magnitude less likely due to the transfer rates decaying exponentially with increasing distance.

The blue band in Fig. 4e shows the time-evolution of *G*(*t*) computed with the rate constants obtained from real-time ET calculations which account for molecular vibrations. The curve shows the same trend as the ground state population derived from the experimental rates. Interestingly, the small backward transfer rate constants obtained from the Marcus theory models 1 (Cascone *et. al*. [41]) and model 2 (Xu *et. al*. [11]) prevent even qualitative agreement with the experimentally predicted dependencies. To evaluate the effect of the chosen value of *k*_*AG*_, we have fitted *k*_*AG*_ and shown that it closely matches the experimental result. The results of the real-time ET calculations match well with the experimental *G*(*t*) curve assuming *k*_*AG*_ = 101 ps. For the Marcus theory estimates, the qualitative disagreement persists even for the best possible value of *k*_*AG*_. In particular, the Marcus theory models with fitted *k*_*AG*_ predict a convergence of *G*(*t*) to a value of 0.4-0.5 within 1-10 ps. Overall, the experimentally observed recombination rates on the order of 10-100 ps are in line with the rates observed for similar proteins harboring an ET chain [32–34].

## Discussion

To summarize, this manuscript discussed several aspects of the light-induced ET in *Er*Cry4a. By comparing Marcus theory estimates with the timescales of structural changes inside of the protein, we laid out how standard Marcus theory neglects the truly non-equilibrium nature of the ET. The energetically excited state induced through the photon absorption as well as the close proximity of nearly indistinguishable electronic states lead to picosecond time-scale ETs, while the slow, globular motions of the protein require hundreds of nanoseconds to fall into a new equilibrium: an observation that disagrees with the equilibrium assumption of Marcus theory. In contrast, real-time QM/MM calculations allowed to discern the coupling between the ET and the relaxation of the protein environment. The crucial role for molecular vibrations in facilitating electron transfers was exemplified via numerical simulations and was analytically rationalized through the analysis of molecular Huang-Rhys factors. The Huang-Rhys factor allowed us to quantify the energy scale of ET induced molecular vibrations in the case of TrpH ionization as 5-7 kcal/mol. The analysis allowed us to bring the interpretation of transient absorption spectra recorded for waiting times of up to 1 ns in close agreement with non-equilibrium ET calculations as demonstrated through the predicted recombination rate shown in Fig. 4e. The closest match between the computationally predicted kinetics and the experimental decay behavior was obtained through assuming the 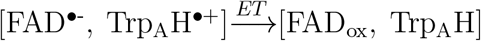 recombination to occur with a time constant of 101 ps, which is in quantitative agreement with the earlier experimental studies on a mutated variant of *Er*Cry4a suggesting a time constant of 89 ps [14].

Our results show that local, non-equilibrium protein and solvent fluctuations play a fundamental role for the ET dynamics of flavoproteins in accordance with previous studies [35, 59]. In extension to those previous studies, we demonstrate here how a highly approximative, atomistic computational approach can reveal the effect of non-equilibrium configurations and vibrational couplings and how they can refine our understanding of experimental observations.

We emphasize that the present study is restricted to the early time dynamics (up to 1 ns) of the radical pairs. It has been demonstrated experimentally [11], that measurable radical pair concentrations, likely formed by deprotonation of TrpH^•+^, persist in cryptochrome for up to 9 *µ*s after photoexcitation [11]. The analysis of such long-lived radical pairs goes beyond the present work. Within the limited timescale of the experiments and simulations, it appears that no more than 40% of the radical pairs that were created through the photo-activation survive for more than 1 ns, representing an upper bound to the quantum yield of the system. Beyond 1 ns, we expect further structural relaxation to be relevant for the stabilization of said radical pairs (see supplementary results). The presented results are in agreement with the existence of long-lived radical pairs if further protein structure relaxation significantly reduces the backward electron transfer rates within 10-100 ns after photo-excitation.

Our observations strongly support the interpretation that ETs in proteins usually are non-equilibrium, non-ergodian processes [36, 38]. Real-time QM/MM calculations appear to be an appropriate method to describe these ETs. The extension of physical realism along with time-dependent descriptions is a possible path for future studies. The inclusion of more accurate QM or MM descriptions, an inclusion of alternative tryptophan transport chains and the extension of simulation timescales would allow us to gain a more reliable picture of the ET within *Er*Cry4a specifically. Alternatively, the simultaneous treatment of spin-dynamics may unravel yet purely neglected physical contributions and might allow to construct a definition of the spin-correlation time within this biological system that incorporates electronic, magnetic and spin interactions in a concerted way.

## Methods

### Sample preparation

Wild-type *Er*Cry4a (GenBank accession KX890129.1) was produced and purified broadly following a previously published protocol [11], with several adjustments. Briefly, His-tagged *Er*Cry4a was heterologously expressed in Escherichia coli BL21(DE3) cells under dark conditions using lysogeny broth supplemented with 10 g/l yeast extract, rather than 5 g/l. The duration of protein expression was increased from 22 h to 44 h. All subsequent purification steps were carried out under dim red light. The recombinant protein was first isolated using Ni-NTA affinity chromatography and further purified by anion exchange chromatography. Purified *Er*Cry4a was concentrated to 5-6 mg/ml in buffer containing 20 mM Tris, 250 mM NaCl, 20% (v/v) glycerol, and 10 mM 2-mercaptoethanol. Protein preparations were snap-frozen in liquid nitrogen and stored at −20 degrees Celcius until further analysis.

### Pump-probe spectroscopy of wildtype *Er*Cry4a

Pump-probe experiments were performed using a setup previously described in Ref. [14] and following the sample handling and experimental details described therein. A titan-sapphire amplifier system (Legend Elite, Coherent) operating at 10 kHz repetition rate and delivering 25 fs and 1 mJ pulses at 800 nm was used to drive an optical parametric amplifier (TOPAS, Light Conversion) in order to generate pump pulses centered at 450 nm with 30 fs pulse duration. A small fraction of the 800 nm fundamental light was used to generate a supercontinuum in CaF_2_ crystal that is continuously moved to avoid photodamage. These pulses were filtered to 320-750 nm and used as the probe pulses. Pump and probe were both focused into the sample cuvette to 50 µm spot size and the transmitted probe was sent to a grating spectrograph (Acton SP2150i, Princeton Instruments) with a mounted fast and sensitive line camera (Aviiva EM4, e2v) that records probe spectra S(*λ*) at full laser repetition rate. A delay *t*_*d*_ between pump and probe is set via a motorized linear translation stage (M531.5IM, Physik Instrumente). An optical chopper system (MC2000, Thorlabs) with a custom-made wheel was used to periodically modulate both pump and probe on a single shot basis to record differential transmission spectra

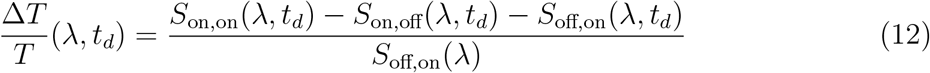

where *S*_(pump,probe)_(*λ*) denotes the pump/probe either being blocked (off) or transmitted (on). This enables dynamic scattering correction [14, 60] helpful for measuring protein samples. A pump pulse energy of 20 nJ and probe pulse energy of 1 nJ was used for which the sample non-linearity was still expected to be in the linear regime [14]. During the experiment, the sample chamber was continuously moved up and down to reduce laser exposure at any given position for prolonged time. No sample degradation was observed during the experiment. A reference measurement on bare buffer solution was recorded under identical conditions to independently measure the cross-phase modulation (XPM) signal [55, 56, 61] that marks the wavelength-dependent zero point reflecting the few-ps probe chirp.

### Analysis of the pump-probe data

Experimental pump-probe maps were recorded for parallel and crossed polarization between pump and probe, averaged and magic angle data computed [14, 55]. These were then chirp- and solvent-corrected [14, 55, 56] using the XPM signal of the buffer reference measurement. After removing data up to *t*_*d*_=200 fs that were contaminated by residual XPM, a global analysis [14, 55, 57] was used to decompose the experimental data into a set of *n* decay-associated difference spectra (*DADS*_*i*_, *i* = 1… *n*) with exponential decay times *τ*_*i*_ such that the total signal can be expressed as

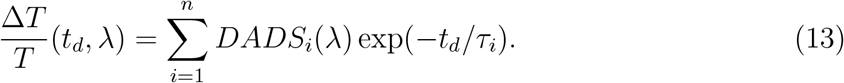

The smallest number of components was determined to be *n* = 5 for which residual spectral difference maps between the experimental data and the fit only show noise around zero without any structure. To compute more physically interpretable spectra, the DADS were transformed into evolution-associated difference spectra

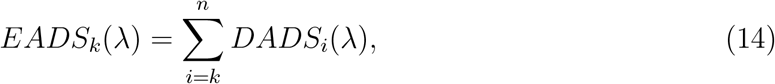

which reflect species spectra of a consecutive rate equation model. The obtained EADS and decay times appeared to be in a good agreement with those previously reported for wildtype *Er*Cry4a [14].

### Classical molecular dynamics simulations

The structure of *Er*Cry4a in its dark state was adapted from earlier studies [8, 11, 43] where it was simulated in a 94 Å×106 Å×102 Å water box, neutralized with 0.15 M NaCl, resulting in a total of 100,518 atoms. The earlier simulations of the structure included 250 ns of production simulation, and was used as the starting structure for the simulations employed in the present study. The structure used from earlier simulations was further simulated utilizing the GROMACS 2022 software [62]. The structure was minimized for 500 conjugate gradient steps using steepest descent minimization algorithm, followed by a second, 2 ns equilibration using the leap-frog integrator with a temperature of 300 K, kept constant by a Berendsen thermostat. The equilibration was followed by a 100 ns dynamic equilibration and 100 ns production simulation. Both equilibration and production simulations utilized a 2 fs timestep and the LINCS algorithm [63] to keep the bond lengths involving hydrogen atoms fixed at their equilibrium values. Periodic boundary conditions were adopted for all stages, and the particle-mesh-Ewald summation method was employed for evaluating the Coulomb forces. Van der Waals forces were calculated using a smooth cut-off distance of 12 Å with a switching distance of 10 Å. The molecular dynamics calculation utilized the Amber99SB forcefields for proteins [64, 65] with earlier parametrized forcefields for the FAD cofactor [47, 48, 66].

### Real-time ET simulations

The method for the real-time electron transfer has been carried out using a hybrid QM/MM DFTB-like protocol, which has been utilized in earlier studies of cryptochromes [14, 47, 67]. The method is outlined and explained in further detail in the earlier articles by Kubar, Elstner *et al*. [47, 48]. The protocol employed molecular fragments participating in the electron transfer as a defined segment of the system that was subjected to the quantum mechanical description (QM region), with the rest of the system being described using classical molecular mechanics force fields (MM region), with the MM region being calculated through usual molecular dynamics methods and the QM region having the kinetic part being described as the derivative of energy with respect to the nuclear coordinates and the electrostatic interactions being calculated by projecting the unpaired electron onto the atomic charges. The QM region is purely used for energetic calculations and will not subject any coordinate changes to the molecular dynamics trajectory, solely introduce the action of the electron transfer itself. The QM region in the present study consists of the conserved tryptophan tetrad [68], and comprises solely of the four tryptophan residues with no linkers needed to facilitate the electron transfer. The QM computations utilizes a separation of frontier orbitals similar to the Hückel and Pariser-Parr-Pople models where the orbitals are assumed to be based on molecular fragments instead of atoms [45, 69]. The DFTB calculations were carried out starting from 50 different structures, sampled every 1 ns from the last 50 ns of the production molecular dynamics simulation.

The tryptophans in the QM region were assumed in the initial state Trp_A_H^•+^. The missing electron on the first tryptophan of the tetrad is assumed to having been transferred to the FAD i.e., being in its FADH^•-^ state. The snapshots were each simulated for 1 ns using a time step of 1 fs. For the fully constrained simulations another set of 50 trajectories was simulated for a shorter simulation time of 630 ps. The MM region was simulated using the Amber99SB forcefields for proteins [64, 65] with earlier parametrized forcefields for the FAD cofactor [47, 48, 66], while the QM region and electron transfer was estimated using earlier parameters [46, 70].

The simulation data with constrained vibrations used the numerical LINCS algorithm to constrain all bond lengths to their equilibrium value. Those simulations have been utilized in a previous study [14]. In the simulations with free vibrations, no bond length constrains were utilized. The constrained vibration simulations consists of 100 QM/MM trajectories initialized from different initial configurations taken from ground state (dark state) molecular dynamics. For the set of freely vibrating simulations, 50 trajectories were simulated.

## Supporting information

Complete_Supplementary_Information

## Data availability

The complete, yet subsampled, trajectories along with the experimental data are accessible here: https://dare.uol.de/privateurl.xhtml?token=87d2f56b-04fe-4eff-a326-e06242be1ed7

This link will be replaced by a DOI, once it is clear that the review and publication process does not require an expansion of the data repository.

## Acknowledgement(s)

IAS thanks the Deutsche Forschungsgemeinschaft (DFG) for its support through TRR386, HYP*MOL, project number 514664767; IAS, CL, HM, KK, ADS and JH thank the DFG for funding through SFB 1372, Magnetoreception and Navigation in Vertebrates, No. 395940726 and EXC-3051, Excellence Cluster NaviSense, No. 533653176. IAS, CL and ADS are grateful to the Ministry of Science and Culture of Lower Saxony (Dynamik auf der Nanoskala: Von kohärenten Elementarprozessen zur Funktionalität (DyNano)). CL and ADS thank the Deutsche Forschungsgemeinschaft Li 580/16-1, DE 3578/3-1) and the Niedersächsische Ministerium für Wissenschaft und Kultur (Wissenschaftsraum ElLiKo). HM, IAS, CL thank “Excellenzstärken”. The authors gratefully acknowledge the computing time granted by the Resource Allocation Board and provided on the supercomputer Emmy/Grete at NHR-Nord@Güttingen as part of the NHR infrastructure. The calculations for this research were conducted with computing resources under the project nip00058.

## Author contributions

IAS and CL conceptualized the study. AF performed simulations. JH and AF analyzed and interpreted simulation data. DTh and JH derived the analytical theory. GS, JS and RB prepared and purified the tested protein samples under supervision of RB and HM. DTi, DL conducted spectroscopy experiments. DTi processed the spectroscopic data. DTi and CL analyzed and interpreted spectroscopic data with help from ADS and KK. IAS, CL, HM and ADS gathered funding and resources. JH and DTi wrote the initial draft and created the figures. All authors reviewed the manuscript.

## References

[1] Martin, D. R. & Matyushov, D. V. Electron-transfer chain in respiratory complex I. Scientific Reports 7, 5495 (2017). URL 10.1038/s41598-017-05779-y.

[2] Richardson, K. H. et al. Functional basis of electron transport within photosynthetic complex I. Nature Communications 12, 5387 (2021). URL 10.1038/s41467-021-25527-1.

[3] Christou, N.-E. et al. Time-resolved crystallography captures light-driven DNA repair. Science 382, 1015–1020 (2023). URL 10.1126/science.adj4270.

[4] Jepsen, K. A. & Solov’yov, I. A. On binding specificity of (6-4) photolyase to a T(6-4)T DNA photoproduct. Europ. Phys. J. D 71, 155–165 (2017). URL https://link.springer.com/article/10.1140/epjd/e2017-70818-2.

[5] Sjulstok, E., Olsen, J. M. H. & Solov’yov, I. A. Quantifying electron transfer reactions in biological systems: what interactions play the major role? Sci. Rep. 5, 18446 (2015). URL 10.1038/srep18446.

[6] Luo, J., Hungerland, J., Solov’yov, I. A., Subotnik, J. E. & Hammes-Schiffer, S. Protein and solvent reorganization drives radical pair stability in avian cryptochrome 4a. Journal of the American Chemical Society 147, 43934–43945 (2025). URL 10.1021/jacs.5c15726.

[7] Ritz, T., Adem, S. & Schulten, K. A model for photoreceptor-based magnetoreception in birds. Biophysical Journal 78, 707–718 (2000). URL 10.1016/s0006-3495(00)76629-x.

[8] Günther, A. et al. Double-cone localization and seasonal expression pattern suggest a role in magnetoreception for european robin cryptochrome 4. Current Biology 28, 211–223.e4 (2018). URL 10.1016/j.cub.2017.12.003.

[9] Mouritsen, H. Long-distance navigation and magnetoreception in migratory animals. Nature 558, 50–59 (2018). URL 10.1038/s41586-018-0176-1.

[10] Hore, P. J. & Mouritsen, H. The radical-pair mechanism of magnetoreception. Annual Review of Biophysics 45, 299–344 (2016). URL 10.1146/annurev-biophys-032116-094545.

[11] Xu, J. et al. Magnetic sensitivity of cryptochrome 4 from a migratory songbird. Nature 594, 535–540 (2021). URL 10.1038/s41586-021-03618-9.

[12] Alvarez, P. H., Gerhards, L., Solov’yov, I. A. & de Oliveira, M. C. Quantum phenomena in biological system. Front. Quantum Sci. Technol. 3, 1466906–(1–13) (2024). URL 10.3389/frqst.2024.1466906.

[13] Pedersen, J. B., Nielsen, C. & Solov’yov, I. A. Multiscale description of avian migration: from chemical compass to behaviour modeling. Sci. Rep. 6, 36709 (2016). URL http://www.nature.com/articles/srep36709.

[14] Timmer, D. et al. Tracking the electron transfer cascade in european robin cryptochrome 4 mutants. Journal of the American Chemical Society 145, 11566–11578 (2023). URL 10.1021/jacs.3c00442.

[15] Langebrake, C. et al. Adaptive evolution and loss of a putative magnetoreceptor in passerines. Proceedings of the Royal Society B: Biological Sciences 291 (2024). URL 10.1098/rspb.2023.2308.

[16] Mouritsen, H., Feenders, G., Liedvogel, M., Wada, K. & Jarvis, E. D. Night-vision brain area in migratory songbirds. Proceedings of the National Academy of Sciences 102, 8339–8344 (2005). URL https://www.pnas.org/doi/abs/10.1073/pnas.0409575102.

[17] Zapka, M. et al. Visual but not trigeminal mediation of magnetic compass information in a migratory bird. Nature 461, 1274–1277 (2009).

[18] Chetverikova, R., Dautaj, G., Schwigon, L., Dedek, K. & Mouritsen, H. Double cones in the avian retina form an oriented mosaic which might facilitate magnetoreception and/or polarized light sensing. Journal of The Royal Society Interface 19, 20210877 (2022). URL 10.1098/rsif.2021.0877.

[19] Wiltschko, R., Stapput, K., Thalau, P. & Wiltschko, W. Directional orientation of birds by the magnetic field under different light conditions. Journal of The Royal Society Interface 7 (2009). URL 10.1098/rsif.2009.0367.focus.

[20] Leberecht, B. et al. Upper bound for broadband radiofrequency field disruption of magnetic compass orientation in night-migratory songbirds. Proceedings of the National Academy of Sciences 120 (2023). URL 10.1073/pnas.2301153120.

[21] Gerhards, L., Deser, A., Kattnig, D. R., Matysik, J. & Solov’yov, I. A. Weak radiofrequency field effects on biological systems mediated through the radical pair mechanism. Chemical Reviews 125, 8051–8088 (2025). URL 10.1021/acs.chemrev.5c00178.

[22] Leberecht, B. et al. Broadband 75–85 MHz radiofrequency fields disrupt magnetic compass orientation in night-migratory songbirds consistent with a flavin-based radical pair magnetoreceptor. J. Comp. Physiol. A 208, 97–106 (2022).

[23] Schwarze, S. et al. Weak broadband electromagnetic fields are more disruptive to magnetic compass orientation in a night-migratory songbird (erithacus rubecula) than strong narrow-band fields. Front. Behav. Neurosci. 10, 55 (2016).

[24] Engels, S. et al. Anthropogenic electromagnetic noise disrupts magnetic compass in a migratory bird. Nature 509, 353–356 (2014).

[25] Ritz, T., Thalau, P., Phillips, J. B., Wiltschko, R. & Wiltschko, W. Resonance effects indicate a radical-pair mechanism for avian magnetic compass. Nature 429, 177–180 (2004).

[26] Kattnig, D. R., Solov’yov, I. A. & Hore, P. J. Electron spin relaxation in cryptochrome-based magnetoreception. Physical Chemistry Chemical Physics 18, 12443–12456 (2016). URL 10.1039/C5CP06731F.

[27] Grüning, G., Gerhards, L., Wong, S. Y., Kattnig, D. R. & Solov’yov, I. A. The effect of spin relaxation on magnetic compass sensitivity in ErCry4a. ChemPhysChem 25, e202400129 (2024). URL https://chemistry-europe.onlinelibrary.wiley.com/doi/abs/10.1002/cphc.202400129.

[28] Gürtemaker, K. et al. Direct interaction of avian cryptochrome 4 with a cone specific G-protein. Cells 11, 2043 (2022). URL https://www.mdpi.com/2073-4409/11/13/2043.

[29] Wu, H., Scholten, A., Einwich, A., Mouritsen, H. & Koch, K. W. Protein-protein interaction of the putative magnetoreceptor cryptochrome 4 expressed in the avian retina. Sci. Rep. 10, 7364 (2020). URL https://www.nature.com/articles/s41598-020-64429-y.

[30] Yee, C. et al. Comparison of retinol binding protein 1 with cone specific g-protein as putative effector molecules in cryptochrome signalling. Scientific Reports 14, 28326 (2024). URL https://www.nature.com/articles/s41598-024-79699-z.

[31] Cellini, A. et al. Directed ultrafast conformational changes accompany electron transfer in a photolyase as resolved by serial crystallography. Nature Chemistry 16, 624–632 (2024). URL 10.1038/s41557-023-01413-9.

[32] Martin, R. et al. Ultrafast flavin photoreduction in an oxidized animal (6-4) photolyase through an unconventional tryptophan tetrad. Physical Chemistry Chemical Physics 19, 24493–24504 (2017). URL 10.1039/c7cp04555g.

[33] Kutta, R. J., Archipowa, N. & Scrutton, N. S. The sacrificial inactivation of the blue-light photosensor cryptochrome from drosophila melanogaster. Physical Chemistry Chemical Physics 20, 28767–28776 (2018). URL 10.1039/C8CP04671A.

[34] Lacombat, F. et al. Ultrafast oxidation of a tyrosine by proton-coupled electron transfer promotes light activation of an animal-like cryptochrome. Journal of the American Chemical Society 141, 13394–13409 (2019). URL 10.1021/jacs.9b03680.

[35] Lu, Y., Kundu, M. & Zhong, D. Effects of nonequilibrium fluctuations on ultrafast short-range electron transfer dynamics. Nature Communications 11, 2822 (2020). URL 10.1038/s41467-020-15535-y.

[36] Matyushov, D. V. Reorganization energy of electron transfer. Physical Chemistry Chemical Physics 25, 7589–7610 (2023). URL 10.1039/D2CP06072H.

[37] Matyushov, D. V. Protein electron transfer: is biology (thermo)dynamic? Journal of Physics: Condensed Matter 27, 473001 (2015). URL 10.1088/0953-8984/27/47/473001.

[38] LeBard, D. N. & Matyushov, D. V. Protein–water electrostatics and principles of bioenergetics. Physical Chemistry Chemical Physics 12, 15335 (2010). URL 10.1039/C0CP01004A.

[39] Blumberger, J. Recent advances in the theory and molecular simulation of biological electron transfer reactions. Chemical Reviews 115, 11191–11238 (2015). URL 10.1021/acs.chemrev.5b00298.

[40] Thoele, D. An Analytical Theory for Vibrationally-Assisted Electron Transfer Between Molecules. Master’s thesis, Carl von Ossietzky Universitaet Oldenburg (2025).

[41] Cascone, M., Mazzeo, P., Cupellini, L. & Mennucci, B. Multiscale simulation of photoinduced electron transfer in cryptochrome 4 from european robin and pigeon indicates a conserved dynamics. The Journal of Physical Chemistry Letters 16, 8877–8884 (2025). URL 10.1021/acs.jpclett.5c01814.

[42] Hungerland, J., Frederiksen, A., Gerhards, L. & Solov’yov, I. A. Studying folding ↔ unfolding dynamics of solvated alanine polypeptides using molecular dynamics. Eur. Phys. J. D 76, 154–(1–13) (2022). URL 10.1140/epjd/s10053-022-00475-7.

[43] Hanić, M. et al. Computational reconstruction and analysis of structural models of avian cryptochrome 4. The Journal of Physical Chemistry B 126, 4623–4635 (2022). URL 10.1021/acs.jpcb.2c00878.

[44] Kubař, T. & Elstner, M. A hybrid approach to simulation of electron transfer in complex molecular systems. Journal of The Royal Society Interface 10, 20130415 (2013). URL 10.1098/rsif.2013.0415.

[45] Elstner, M. et al. Self-consistent-charge density-functional tight-binding method for simulations of complex materials properties. Physical Review B 58, 7260–7268 (1998). URL 10.1103/PhysRevB.58.7260.

[46] Kubař, T. & Elstner, M. Efficient algorithms for the simulation of non-adiabatic electron transfer in complex molecular systems: application to dna. Physical Chemistry Chemical Physics 15, 5794 (2013). URL 10.1039/C3CP44619K.

[47] Sjulstok, E., Lüdemann, G., Kubař, T., Elstner, M. & Solov’yov, I. A. Molecular insights into variable electron transfer in amphibian cryptochrome. Biophys. J. 114, 2563–2572 (2018). URL https://www.cell.com/biophysj/fulltext/S0006-3495(18)30459-4.

[48] Lüdemann, G., Solov’yov, I. A., Kubař, T. & Elstner, M. Solvent driving force ensures fast formation of a persistent and well-separated radical pair in plant cryptochrome. J. Am. Chem. Soc. 137, 1147–1156 (2015). URL http://pubs.acs.org/doi/abs/10.1021/ja510550g.

[49] Solov’yov, I. A.Chang, P.-Y. & Schulten, K. Vibrationally assisted electron transfer mechanism of olfaction: myth or reality? Physical Chemistry Chemical Physics 14, 13861 (2012). URL 10.1039/C2CP41436H.

[50] Wei, Y.-C. & Hsu, L.-Y. Polaritonic Huang-Rhys factor: Basic concepts and quantifying light–matter interactions in media. The Journal of Physical Chemistry Letters 14, 2395–2401 (2023). URL 10.1021/acs.jpclett.3c00065.

[51] Hsu, C.-P. Reorganization energies and spectral densities for electron transfer problems in charge transport materials. Physical Chemistry Chemical Physics 22, 21630–21641 (2020). URL 10.1039/D0CP02994G.

[52] Zhao, Y. & Truhlar, D. G. The M06 suite of density functionals for main group thermo-chemistry, thermochemical kinetics, noncovalent interactions, excited states, and transition elements: two new functionals and systematic testing of four m06-class functionals and 12 other functionals. Theoretical Chemistry Accounts 120, 215–241 (2007). URL 10.1007/s00214-007-0310-x.

[53] Neese, F. Software update: the ORCA program system, version 6.0. WIRES Comput. Molec. Sci. 15, e70019 (2025).

[54] Neese, F. Approximate second-order SCF convergence for spin unrestricted wavefunctions. Chem. Phys. Lett. 325, 93–98 (2000).

[55] Timmer, D. et al. Structural flexibility slows down charge transfers in diaminoterephthalate-C60 dyads. The Journal of Physical Chemistry C 128, 2380–2391 (2024). URL 10.1021/acs.jpcc.3c08270.

[56] Timmer, D. et al. Direct observation of Franck–Condon stimulated emission and sub-20 fs relaxation in photoexcited flavins. The Journal of Physical Chemistry Letters 17 (2026). URL 10.1021/acs.jpclett.6c00294.

[57] Rabe, M. Spectram: A MATLAB® and gnu octave toolbox for transition model guided deconvolution of dynamic spectroscopic data. Journal of Open Research Software 8, 13 (2020). URL 10.5334/jors.323.

[58] Schwinn, K., Ferré, N. & Huix-Rotllant, M. UV-visible absorption spectrum of FAD and its reduced forms embedded in a cryptochrome protein. Physical Chemistry Chemical Physics 22, 12447–12455 (2020). URL 10.1039/D0CP01714K.

[59] Kundu, M., He, T.-F., Lu, Y., Wang, L. & Zhong, D. Short-range electron transfer in reduced flavodoxin: Ultrafast nonequilibrium dynamics coupled with protein fluctuations. The Journal of Physical Chemistry Letters 9, 2782–2790 (2018). URL 10.1021/acs.jpclett.8b00882.

[60] Timmer, D., Lünemann, D. C., Riese, S., Sio, A. D. & Lienau, C. Full visible range two-dimensional electronic spectroscopy with high time resolution. Optics Express 32, 835 (2023). URL 10.1364/OE.511906.

[61] Ekvall, K. et al. Cross phase modulation artifact in liquid phase transient absorption spectroscopy. Journal of Applied Physics 87, 2340–2352 (2000). URL 10.1063/1.372185.

[62] Hess, B., Kutzner, C., van der Spoel, D. & Lindahl, E. GROMACS 4: Algorithms for highly efficient, load-balanced, and scalable molecular simulation. Journal of Chemical Theory and Computation 4, 435–447 (2008). URL 10.1021/ct700301q.

[63] Hess, B., Bekker, H., Berendsen, H. J. C. & Fraaije, J. G. E. M. LINCS: A linear constraint solver for molecular simulations. Journal of Computational Chemistry 18, 1463–1472 (1997). URL 10.1002/(SICI)1096-987X(199709)18:12<1463::AID-JCC4>3.0.CO;2-H.

[64] Wang, J., Wolf, R. M., Caldwell, J. W., Kollman, P. A. & Case, D. A. Development and testing of a general amber force field. Journal of Computational Chemistry 25, 1157–1174 (2004). URL 10.1002/jcc.20035.

[65] Lindorff-Larsen, K. et al. Improved side-chain torsion potentials for the Amber ff99SB protein force field. Proteins: Structure, Function, and Bioinformatics 78, 1950–1958 (2010). URL 10.1002/prot.22711.

[66] Solov’yov, I. A., Domratcheva, T., Moughal Shahi, A. R. & Schulten, K. Decrypting cryptochrome: Revealing the molecular identity of the photoactivation reaction. Journal of the American Chemical Society 134, 18046–18052 (2012). URL 10.1021/ja3074819.

[67] Frederiksen, A., Aldag, M., Solov’yov, I. A. & Gerhards, L. Activation of cryptochrome 4 from atlantic herring. Biology 13, 262 (2024). URL 10.3390/biology13040262.

[68] Frederiksen, A. et al. Mutational study of the tryptophan tetrad important for electron transfer in european robin cryptochrome 4a. ACS Omega 8, 26425–26436 (2023). URL 10.1021/acsomega.3c02963.

[69] Kitaura, K., Ikeo, E., Asada, T., Nakano, T. & Uebayasi, M. Fragment molecular orbital method: an approximate computational method for large molecules. Chemical Physics Letters 313, 701–706 (1999). URL 10.1016/S0009-2614(99)00874-X.

[70] Kubař, T. & Elstner, M. Coarse-grained time-dependent density functional simulation of charge transfer in complex systems: Application to hole transfer in DNA. The Journal of Physical Chemistry B 114, 11221–11240 (2010). URL 10.1021/jp102814p.

