## Supplementary material for "Non-equilibrium vibrational dynamics govern ultrafast electron transfer in avian cryptochrome": Complete_Supplementary_Information

May 25, 2026

[1]Institute of Physics, Carl von Ossietzky Universität Oldenburg, Carl-von-Ossietzky-Straße 9-11, 26129 Oldenburg, Germany

[2]Department of Chemistry, School of Engineering Sciences in Chemistry, Biotechnology and Health (CBH), KTH Royal Institute of Technology, Stockholm, Sweden

[3]Institute of Biology and Environmental Sciences, Carl von Ossietzky Universität Oldenburg, Carl-von-Ossietzky-Straße 9-11, 26129 Oldenburg, Germany

[4]Research Centre for Neurosensory Science, Carl von Ossietzky Universität Oldenburg, Carl-von-Ossietzky-Straße 9-11, 26129 Oldenburg, Germany

[5]Center for Nanoscale Dynamics (CENAD), Carl von Ossietzky Universität Oldenburg, Carl-von-Ossietzky-Straße 9-11, 26129 Oldenburg, Germany

‡
†
⋆

### Supplementary results

#### Dependence of electron transfer induced vibrations on the quantum chemical methodology

In Fig. 3c of the main manuscript, the expected vibrational energy of a tryptophan sidechain is shown if an electron was instantaneously moved, for example via an electron transfer to a neighbouring tryptophan. For those results, some gas phase calculations of a tryptophan sidechain were performed to obtain optimized geometries and vibrations of the neutral closed shell and positive radical state. The vibrations were calculated in internal coordinates and the internal Hessian was then used to evaluate Equations 4 and 7 of the main manuscript. Table S1 shows the results from calculations with different DFT functionals. Among all tested functionals, the summed vibrational energy of bonds and angles deviates by more than 2 kcal/mol. However, the vibrational energy is significant compared to the expected thermodynamic noise independent of the tested functionals. Independent of the particular choice of functional, it is expected that the energy transfer from the environment's electrostatics into molecular vibrations represents a non-negligible constituent of the electron transfer process.

Table S1: Huang-Rhys factor based estimate of the vibrational energy of TrpH induced by the removal of an electron (see Fig. 3c of the main manuscript) for different DFT methods all employing the def2-TZVPD basis set.

| Functional | Bond vibrational energy (kcal/mol) | Angular vibrational energy (kcal/mol) | Sum (kcal/mol) |
|---|---|---|---|
| B3LYP | 3.89 | 0.93 | 4.82 |
| $\omega$B97 | 5.82 | 1.34 | 7.16 |
| $\omega$B97M-V | 5.28 | 1.23 | 6.51 |
| M06-2X | 4.88 | 1.14 | 6.02 |

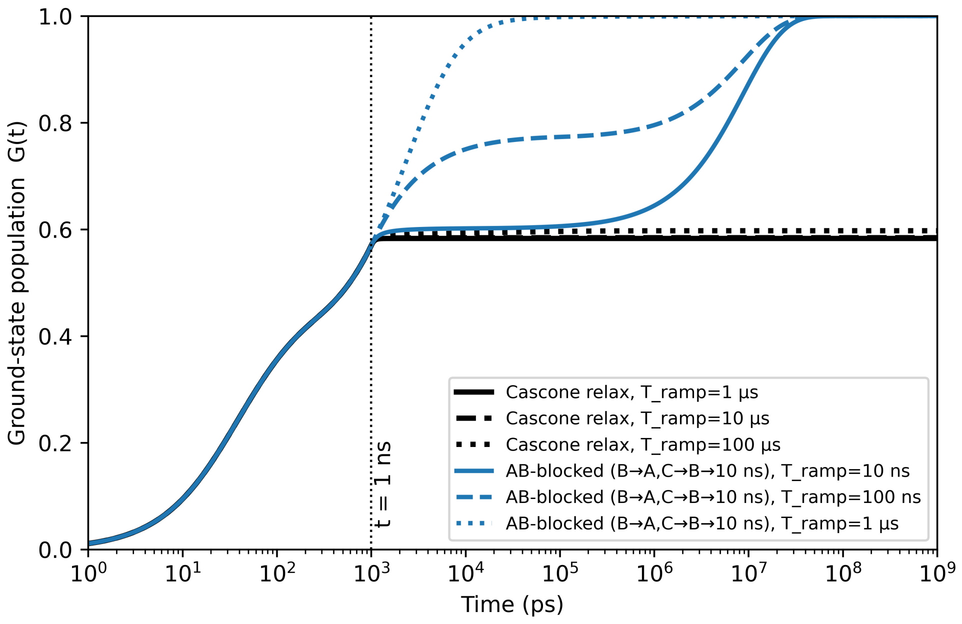


Figure S1: Cumulative recombination probability $G(t)$ extrapolated beyond 1 ns after photoexcitation for different protein relaxation schemes. $G(t)$ is modeled via Eqs. (S1) and (S2). For details on the settings see text.

#### The potential effects of further protein relaxation

The non-equilibrium nature of the forward and backward electron transfers becomes apparent and is discussed in the main manuscript. It is apparent that, based on the forward and backward rates extracted from the numerical simulations for time delays of up to 1 ns, all radical pairs would recombine within 10 ns. This is in disagreement with the experimental observation that long-lived radical pairs persist on the $\mu$s timescale [1]. This suggests that the calculated rates may change at longer timescales due to further protein relaxation.

The equilibration process of the protein gradually establishes the electrostatic potential as illustrated by the electrostatic potentials shown in Fig. 2b of the main manuscript. From the ensemble of simulations, a fixed set of rate constants $k$ was obtained. During further protein equilibration, those rate constants may themselves be subject to change. In that case, the state vector $\vec{s}(t)$ introduced in Eq. 2 of the main manuscript evolves with

$$\frac{d}{dt}\vec{s}(t) = K(t)\vec{s}(t), \tag{S1}$$

such that the coupling matrix becomes time-dependent. Denote the rate constants obtained from the 1 ns long QM/MM trajectories as $K_{\text{initial}}$ and a final set of rate constants as $K_{\text{final}}$. One can then express the time-dependence of the rate constant matrix as

$$K(t) = \begin{cases} K_{\text{initial}} & \text{if } t \leq 1 \text{ ns} \\ \left(1 - \frac{t-1\text{ns}}{T_{\text{ramp}}}\right) K_{\text{initial}} + \frac{t-1\text{ ns}}{T_{\text{ramp}}} K_{\text{final}} & \text{if } 1 \text{ ns} \leq t \leq 1 \text{ ns} + T_{\text{ramp}} \\ K_{\text{final}} & \text{if } 1 \text{ ns} + T_{\text{ramp}} \leq t \end{cases} \tag{S2}$$

Here, $T_{\text{ramp}}$ is the time it takes for the protein to adopt an equilibrium configuration which has the characteristic transfer rates $K_{\text{final}}$.

Figure S1 shows the extrapolated ground state population $G(t)$ for different sets of $K_{\text{final}}$ and $T_{\text{ramp}}$ assuming that the recombination rate from state A, $k_{AG}$ remains constant. For the black set of curves, rate constants from Cascone *et. a.* are used. Even at very slow structural relaxation of $T_{\text{ramp}} = 100$ $\mu$s, recombination stops immediately. The blue curves represent the situation in which only the backward electron transfer rates $k_{BA}$ and $k_{CB}$ are changed and adopt the characteristic time constant of 10 ns. These backward transfer rates are still faster than predicted by the Marcus theory equilibrium estimates shown in Tab. 1 of the main manuscript. Nevertheless, such a modification of the rates would retain 40% of the radical pair for up to 1 $\mu$s in the case of fast relaxation with $T_{\text{ramp}} = 10$ ns or 20% of the radical pair for a more typical relaxation time of $T_{\text{ramp}} = 100$ ns. For comparison, the modulation frequency of the geomagnetic field on an electron spin system is about 700 ns [2].

The above outlined model considerations are by no means a theoretical prediction for the survival probability of long-lived radical pairs in *Er*Cry4a. Within the scope of the

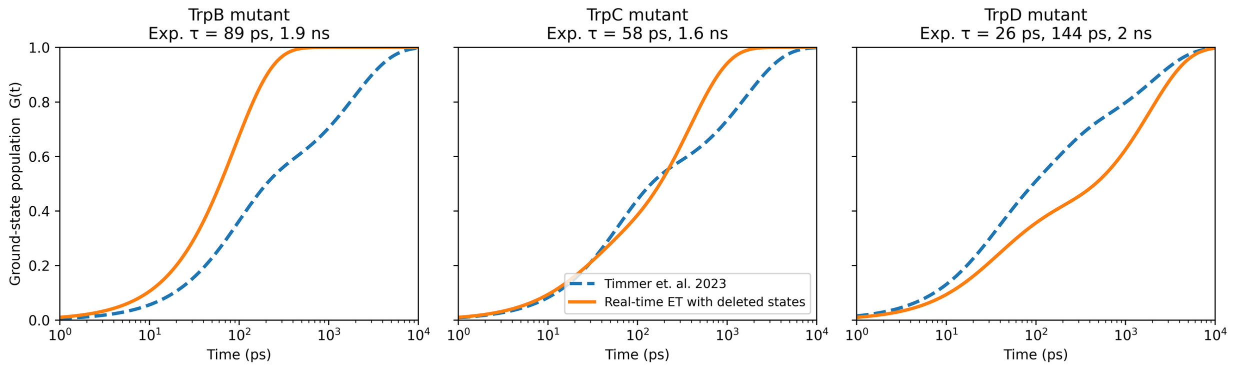


Figure S2: Cumulative recombination probability $G(t)$ for data from mutants obtained by Timmer *et. al.* [3] (blue, dashed lines), compared with theoretical curves that approximate a truncation of the ET chain due to the mutation (orange, solid lines).

present study, it can only be estimated that less than 40% of photo-activations can create a radical pair that survives longer than 1 ns. However, the above models that include a time-dependent set of rates $K(t)$ illustrate, how further equilibration may explain the apparent discrepancy between the here calculated rates and the experimental observation of long-lived radical pairs.

#### Comparison to results from mutated protein variants

A previous study [3] included measurements conducted on mutated variants of $Er$Cry4a. Except for the TrpA mutant, where the ET was interpreted to occur with the adenine instead of the usual ET transfer chain, the mutations of TrpB, TrpC and TrpD were interpreted to truncate the ET chain. To compare the calculated rates with data from those mutated protein variants, one may similarly consider a truncated set of states. Thus, to model the expected curve for $G(t)$ for the TrpB mutant, the state vector only consists of the states $(G, A)$. In the TrpC mutant, states $(G, A, B)$ are included and for the TrpD mutant, states $(G, A, B, C)$ are considered. This can be modeled by simply setting all rate constants in the matrix $K$ that involve the disallowed states to 0.

Figure S2 compares the time-dependence of the recombined population $G(t)$ between experiment (blue, dashed lines) and the truncated rate models (orange, solid lines). For the TrpB mutant data, the theoretical model predicts significantly faster recombination by roughly two orders of magnitude. A possible reason may be the involvement of alternative ET pathways or the occurrence of direct [$FAD^{\bullet -}$, $Trp_A H^{\bullet +}$]$\xrightarrow{ET}$[$FAD^{\bullet -}$, $Trp_C H^{\bullet +}$] transfers. Even though a truncated rate model without any adjustment on the rate constants is a crude approximation, the TrpC and TrpD mutant models show qualitative agreement with the experimental curves.

#### Comparison of the molecular Huang-Rhys factor $\sigma_{n,k}$ with other descriptions of electron-vibrational coupling

The presented Huang-Rhys factor extends the theory introduced by Solov'yov *et. al.* [4] to arbitrary initial and final vibrational states. Other formulations exist that similarly reference the name of a Huang-Rhys factor, for example in the context of an excited solid state system [5]. For a single mode, Wei *et. al.* [5] utilize the vibronic Huang-Rhys factor

$$S_{vib} = \frac{m\omega_{vib}\Delta X^2}{2\hbar}. \tag{S3}$$

Here, $m$ is the mass associated with a vibrational mode with frequency $\omega_{vib}$, $\hbar$ is the reduced Planck's constant and $\Delta X$ is the displacement. To showcase the similarity to the molecular Huang-Rhys factor $\sigma_{n,k}(u)$, consider the coupling between an initial ground state, $k = 0$, with the first excited state after the transfer, $n = 1$. Equation (4) of the main manuscript reduces to

$$\sigma_{1,0}(u) = \left| \frac{(-1)ue^{-u^2/2}}{1} \right|^2 = u^2 e^{-u^2}. \tag{S4}$$

By using the definition of $u$ and approximating $\exp(-u^2) \approx 1$ for small displacements $q$, one arrives at

$$\sigma_{1,0}(q) \approx \frac{m\omega q^2}{2\hbar}. \tag{S5}$$

Since $q$ and $\Delta X$ are two different notations for the displacement, the presented *molecular* Huang-Rhys factor can be reduced to the *vibronic* Huang-Rhys factor used by Wei *et. al.* via some additional approximations.

Historically, the Huang-Rhys factor has first been introduced for ionic centers in a solid state system [6]. The term of a Huang-Rhys parameter can be found to describe electron-phonon couplings [7] or electron-polariton couplings [5]. Phonons and polaritons are the quasiparticles for discrete vibrational or polar excitations within a periodic system. In contrast, the *molecular* Huang-Rhys factor discussed in the main manuscript specifically describes a finite, non-periodic system. While the mathematical descriptions and derivations for periodic, infinite systems fundamentally differ from those required for non-periodic, finite systems, the physical description of Huang-Rhys factor follows a common theme. Huang-Rhys factors describe the coupling between electronic states and excitations of vibrating degrees of freedom. In the case of phonons or molecular eigenfrequencies, the vibrations are exerted by nuclear motion.

The displaced harmonic oscillator model for the calculation of reorganization energies [8, 9] is the closest existing analog of the molecular Huang-Rhys factor. In fact, the displaced harmonic oscillator model is recovered from the molecular Huang-Rhys factor via the additional approximations that resulted in Eq. (S5).

### References


[1] Xu, J. *et al.* Magnetic sensitivity of cryptochrome 4 from a migratory songbird. *Nature* **594**, 535–540 (2021). URL `http://dx.doi.org/10.1038/s41586-021-03618-9`.

[2] Hore, P. J. & Mouritsen, H. The radical-pair mechanism of magnetoreception. *Annual Review of Biophysics* **45**, 299–344 (2016). URL `http://dx.doi.org/10.1146/annurev-biophys-032116-094545`.

[3] Timmer, D. *et al.* Tracking the electron transfer cascade in european robin cryptochrome 4 mutants. *Journal of the American Chemical Society* **145**, 11566–11578 (2023). URL `http://dx.doi.org/10.1021/jacs.3c00442`.

[4] Solov'yov, I. A., Domratcheva, T., Moughal Shahi, A. R. & Schulten, K. Decrypting cryptochrome: Revealing the molecular identity of the photoactivation reaction. *Journal of the American Chemical Society* **134**, 18046–18052 (2012). URL `http://dx.doi.org/10.1021/ja3074819`.

[5] Wei, Y.-C. & Hsu, L.-Y. Polaritonic Huang-Rhys factor: Basic concepts and quantifying light–matter interactions in media. *The Journal of Physical Chemistry Letters* **14**, 2395–2401 (2023). URL `http://dx.doi.org/10.1021/acs.jpclett.3c00065`.

[6] Huang, K. & Rhys, A. Theory of light absorption and non-radiative transitions in f - centres. *Proceedings of the Royal Society of London. Series A. Mathematical and Physical Sciences* **204**, 406–423 (1950). URL `http://dx.doi.org/10.1098/rspa.1950.0184`.

[7] Zhang, Y. Applications of huang–rhys theory in semiconductor optical spectroscopy. *Journal of Semiconductors* **40**, 091102 (2019). URL `http://dx.doi.org/10.1088/1674-4926/40/9/091102`.

[8] Hsu, C.-P. Reorganization energies and spectral densities for electron transfer problems in charge transport materials. *Physical Chemistry Chemical Physics* **22**, 21630–21641 (2020). URL http://dx.doi.org/10.1039/D0CP02994G.

[9] Avila Ferrer, F. J. & Santoro, F. Comparison of vertical and adiabatic harmonic approaches for the calculation of the vibrational structure of electronic spectra. *Physical Chemistry Chemical Physics* **14**, 13549 (2012). URL http://dx.doi.org/10.1039/C2CP41169E.
